# The SARS-CoV-2 Nucleocapsid phosphoprotein forms mutually exclusive condensates with RNA and the membrane-associated M protein

**DOI:** 10.1101/2020.07.30.228023

**Authors:** Shan Lu, Qiaozhen Ye, Digvijay Singh, Elizabeth Villa, Don W. Cleveland, Kevin D. Corbett

## Abstract

The multifunctional nucleocapsid (N) protein in SARS-CoV-2 binds the ~30 kb viral RNA genome to aid its packaging into the 80-90 nm membrane-enveloped virion. The N protein is composed of N-terminal RNA-binding and C-terminal dimerization domains that are flanked by three intrinsically disordered regions. Here we demonstrate that a centrally located 40 amino acid intrinsically disordered domain drives phase separation of N protein when bound to RNA, with the morphology of the resulting condensates affected by inclusion in the RNA of the putative SARS-CoV-2 packaging signal. The SARS-CoV-2 M protein, normally embedded in the virion membrane with its C-terminus extending into the virion core, independently induces N protein phase separation that is dependent on the N protein’s C-terminal dimerization domain and disordered region. Three-component mixtures of N+M+RNA form condensates with mutually exclusive compartments containing N+M or N+RNA, including spherical annular structures in which the M protein coats the outside of an N+RNA condensate. These findings support a model in which phase separation of the N protein with both the viral genomic RNA and the SARS-CoV-2 M protein facilitates RNA packaging and virion assembly.

## Introduction

The ongoing COVID-19 pandemic is caused by Severe Acute Respiratory Syndrome Coronavirus 2 (SARS-CoV-2), a highly contagious betacoronavirus (1, 2). Coronaviruses comprise a large family of positive-stranded RNA viruses, whose ~30 kb genome is packaged into a membrane-enveloped virion 80-90 nm in diameter (3, 4). The first two-thirds of the genome encodes two polyproteins that are processed by virally-encoded proteases into nonstructural proteins and which assemble into the viral replicase-transcriptase complex (RTC) (5). The final third of the genome generates sub-genomic RNAs encoding accessory proteins plus the four main structural proteins of the virion: the spike (S) protein that recognizes cell receptors, the nucleocapsid (N) protein responsible for viral RNA packaging, and the membrane-associated envelope (E) and membrane (M) proteins ((6, 7).

The RNA-binding N protein plays two major roles in the coronavirus life cycle. Its primary role is to assemble with genomic RNA into the viral RNA-protein(vRNP) complex, and mediate vRNP packaging into virions via poorly-understood interactions between the N and M (membrane) proteins (8–10). Second, the N protein localizes to replicase-transcriptase complexes at early stages of infection, where it is thought to facilitate viral RNA synthesis and translation by recruiting host factors and promoting RNA template switching (11–15). To accomplish these important functions, betacoronavirus N proteins have evolved a modular architecture with two conserved, folded domains flanked by three intrinsically disordered regions (IDRs). The N-terminal domain (NTD) is thought to mediate a specific interaction with the viral genome’s “packaging signal”, and the C-terminal domain (CTD) forms a compact dimer that has been proposed to aid vRNP assembly (16–21). These two domains are separated by a conserved central IDR containing a serine/arginine-rich region (SR) that is highly phosphorylated in infected cells (22–26), and are flanked by less well-conserved IDRs at the N- and C-terminus. The C-terminal IDR of the related SARS-CoV N protein has been implicated in M protein binding, suggesting an important role for this domain in viral packaging (9, 10).

In isolation, betacoronavirus N proteins self-associate into dimers, tetramers, and larger oligomers that are thought to form the basis for assembly of the vRNP complex (21, 27–29). Electron microscopy analysis of several betacoronaviruses has suggested that in the virion, the N protein mediates assembly of a helical filament-like vRNP complex (3, 30–32). Recent cryo-electron tomography of SARS-CoV-2 virions has revealed a more granular vRNP structure, with each virion containing 35-40 individual vRNP complexes that adopt a cylindrical shell-like structure ~15nm in diameter (4, 33). Many such vRNPs are membrane-proximal and show a characteristic orientation with respect to the membrane, suggesting potential specific interactions with the membrane-associated M protein (33). Within the vRNP, tentative modeling based on known NTD and CTD structures suggests a specific assembly with ~800 nt of genomic RNA (30 kb divided among ~38 vRNPs) wrapped around ~12 copies of the N protein (33). Individual vRNPs could also form linear stacks resembling helical filaments (4, 33), reconciling the apparent conflict with earlier observations and suggesting that betacoronavirus vRNPs may adopt a broadly conserved architecture.

In recent years, many RNA-binding proteins with intrinsically disordered regions have been found to undergo liquidliquid phase separation with RNA, and these biomolecular condensates are thought to orchestrate a large number of important biological processes and in some cases drive disease (34–41). The structural features of the betacoronavirus N protein and its diverse roles in viral RNA metabolism and virion assembly make it tempting to hypothesize that phase separation may play a role in this protein’s functions. Here we combine *in* vitroreconstitution approaches and cellular assays to determine the phase separation behavior of the SARS-CoV-2 N protein. We first demonstrate that RNA can induce the assembly of the N protein into gel-like phase-separated condensates in *vitro,* and pinpoint a ~40-residue region within the central IDR as necessary for RNA-driven phase separation. We also characterized the interaction between N and a soluble fragment of the M protein, which we find independently mediates N protein phase separation through an interaction with the protein’s C-terminal domains. Three-component mixtures of N, M, and RNA are found to assemble into condensates with distinct internal compartments or layers containing N+RNA or N+M, suggesting that M and RNA repel one another despite each interacting with N. This mutual exclusion often yielded two-layered condensates with an internal N+RNA compartment surrounded by an outer layer of N+M. These data provide mechanistic insights into the phase separation behavior of SARS-CoV-2 N, and suggest that virion assembly may be driven by specific but mutually exclusive interactions of N with genomic RNA and the viral membrane-associated M protein.

## Results

### SARS-CoV-2 N undergoes RNA-dependent phase separation

To investigate the mechanistic basis for nucleocapsid-mediated RNA packaging, we first purified bacterially-expressed recombinant full-length N protein. While purification in buffers with high salt concentration (1M NaCl) yielded pure protein, purification in lower-salt buffers resulted in retention of bacterial nucleic acids and aggregation of the protein as judged by size-exclusion chromatography (21). Recognizing this and recent findings that intrinsically-disordered RNA-binding proteins undergo liquid-liquid phase separation in *vitro* (34–41), we investigated whether the SARS-CoV-2 N protein shares this property. We mixed Cy5-labeled full-length N protein with a non-specific 17-mer ssRNA labeled with fluorescein (6-FAM), and observed formation of spherical phase-separated condensates containing both components (Fig. 1A). Condensates formed in a specific range of RNA concentration, above which phase separation was inhibited (Fig. 1A). This “re-entrant” phase separation behavior is commonly observed in two-component systems including RNA-protein mixtures (42). With 10 μM N protein, maximal phase separation occurred in the presence of 5 μM 17-mer RNA (85 μM nucleotide), above which phase separation was inhibited (Fig. 1A). The observation that N forms condensates with a very short RNA suggests that phase separation in this case is driven largely by multivalent protein-protein interactions. In agreement with this idea, purified N formed condensates without addition of RNA (Fig. S1A), but these were much smaller and fewer than those formed in the presence of RNA. We next examined the behavior of the N protein with three larger RNAs derived from the SARS-CoV-2 genome (Fig 1B, S1B, Table S1): (1) UTR265, the first 265 bases of the viral genome which contains the leader sequence (located at nt 20-81) that in mouse hepatitis virus (MHV) was shown to bind the N protein with high affinity (43, 44); (2) UTR1000, the first 1000 bases of the viral genome that includes the 5’-UTR and part of the ORF1ab gene; (3) PS576, a 576-nt sequence located near the end of the ORF1ab gene (nt 19786-20361) corresponding to the putative packaging signal for SARS-CoV viral RNA (45, 46). We found that addition of 20 ng/μL (~62μM nucleotide) viral RNA fragments containing the 5’ UTR to 10 μM N protein induced the formation of round condensates (Fig. 1C). In contrast, addition of the PS576 fragment caused the formation of more amorphous structures resembling a fibrillar network (Fig. 1C). The distinct structures formed with PS576 compared to 17-mer RNA or UTR-derived RNAs suggests that the viral packaging signal may interact differently with the N protein than nonspecific RNA (see Discussion). All of these condensates grew with time and displayed classical behaviors such as droplet fusion (Fig. 1D, S1C). Similarly, N protein and viral RNA also display “re-entrant” phase separation behavior (Fig. S1D). Cryo-electron tomography of N protein+RNA condensates revealed well-defined texture suggestive of internal order (Fig. 1E), which was also observed by super-resolution light microscopy (Fig. S1E). In particular, these condensates resemble electron-dense virus-like particles observed in the periphery of infected cells and also released alongside intact virions, which may comprise packaged RNAs that have not undergone membrane envelopment (33).

**Fig. 1.**
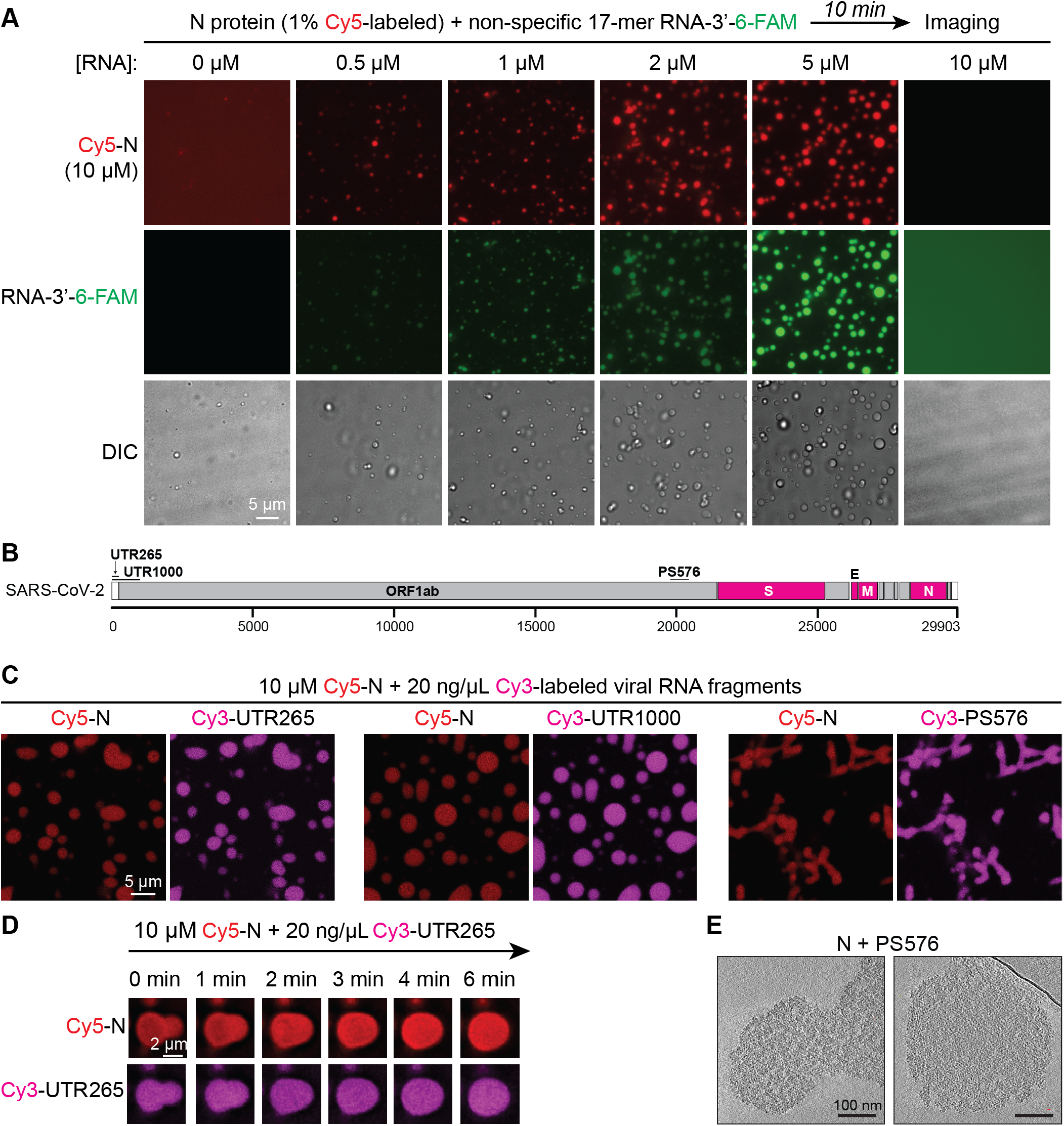
N protein undergoes phase separation with RNA. (A) Fluorescence and DIC images of phase separation of N protein (1% N-terminal Cy5-labeled N protein) with a 3’ 6-FAM labeled 17-mer ssRNA. Scale bar, 5 μm. See Fig. S1A for N protein phase separation in the absence of added RNA. (B) Schematic of SARS-COV-2 viral genome, showing the locations of four structural protein genes (pink) and three viral genome fragments used in this study (Table S1). (C) Fluorescence images of phase separated condensates formed by N protein with viral RNA fragments. Scale bar, 10 μm. (D) Fluorescence images of a representative fusion event of N+RNA condensates. Events were observed in 20 min after initial mixing. (E) Cryo-electron tomography of N protein+RNA condensates. Mixtures of 10 μM N protein and 20 ng/μL PS576 were immediately plunged onto grids. Scale bar, 100 nm.

We also determined the effect of salt concentration on the formation of gel-like condensates with N protein. The N protein formed condensates on its own at very low salt concentrations (20 mM KCl), while addition of RNA enhanced N protein phase separation at salt concentrations approaching physiological (higher than 80 mM KCl) (Fig. S1F). Thus, while the N protein has an inherent tendency to de-mix through self-interactions, additional interactions with RNA (or other proteins; see below) enhance phase separation in physiologically-relevant conditions.

Next, we explored the biophysical properties of N+RNA condensates by fluorescence recovery after photo-bleaching (FRAP). Full bleaching and partial bleaching of N+RNA condensates indicated that the N protein is very slowly exchanged with the soluble pool and within the structures, with recovery of only ~12% fluorescence intensity in 2 minutes after bleaching (Fig. 2). In condensates formed with the short 17-mer RNA, the RNA was highly mobile and freely exchanged (Fig. 2A-B). The mobility of this short RNA decreased with longer incubation times, indicating that initial liquid-like condensates mature to a more solid gel-like structure with time (Fig. S2A). In condensates of N protein with the larger virus-derived RNAs, both RNA and N protein showed slow dynamics (Fig. 2C-D). N protein dynamics were independent of the length of RNA in the condensates (Fig. 2C-D), but in these condensates the longer RNAs showed much slower dynamics than the N protein, with the longest viral RNA showing the lowest mobility. These data suggest that long viral RNAs interact with multiple N proteins and are trapped inside a framework formed by the N protein. Consistent with these slow dynamics, the fusion of N protein droplets was slow (Fig. 1D) and soluble N protein was incorporated very slowly into pre-existing N+RNA condensates (Fig. S2B).

**Fig. 2.**
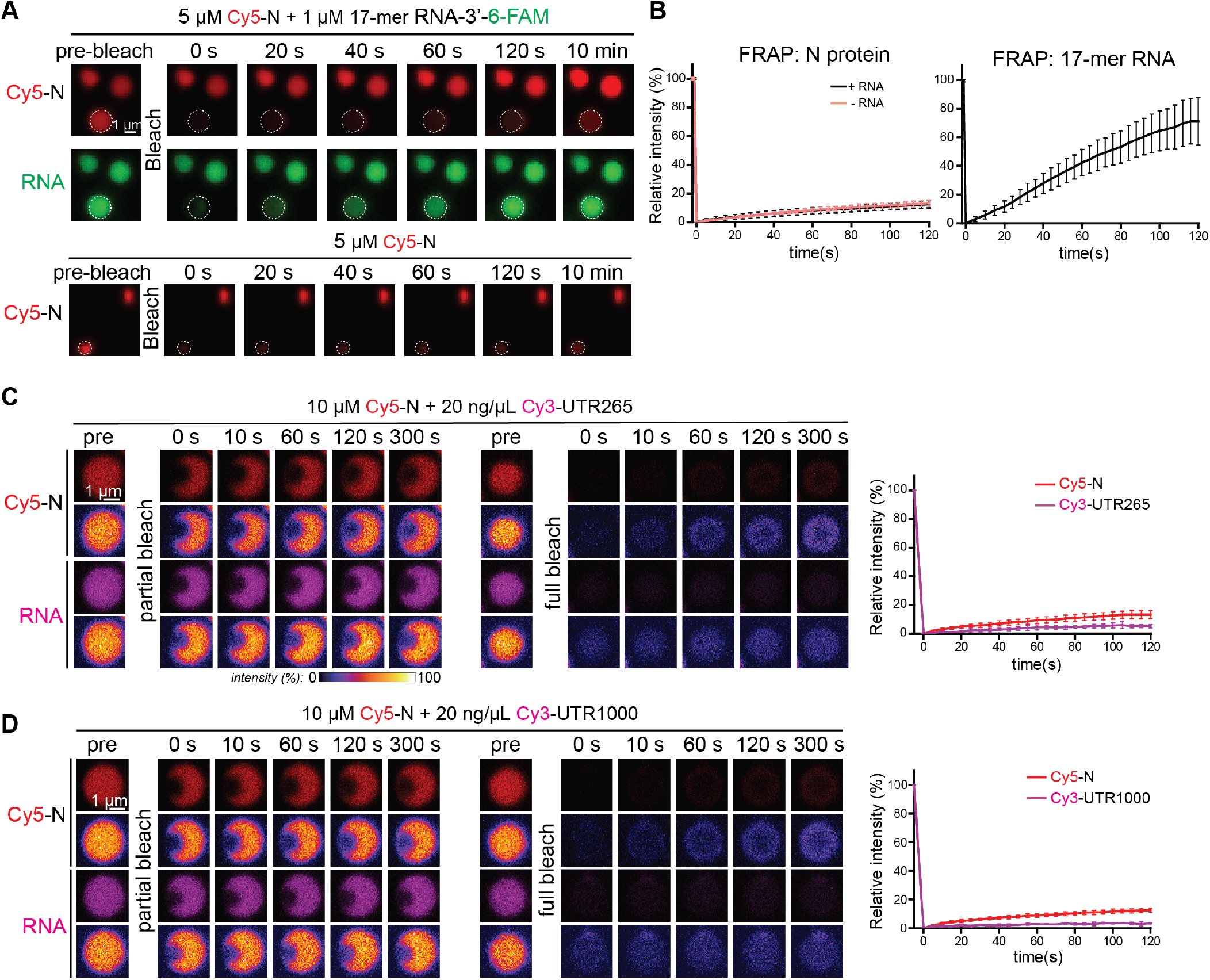
N+RNA forms gel-like structures with slow dynamics. (A-B) FRAP analysis of condensates formed by N protein and short 17-mer ssRNA or N protein only. Representative fluorescence images (A) and mean fluorescence plots (B) of FRAP of N protein + 17-mer ssRNA condensates (n=2) or N protein only condensates (n=4). Data are normalized to the average intensity of a particle before photobleaching and are represented as mean ± standard deviation from the recovery curves. Scale bar, 1 μm. (C-D) FRAP analysis of condensates formed by N protein and viral RNAs: (C) UTR265 (n=8); (D) UTR1000 (n=8). Fluorescence images of one partial-bleached RNA/N protein condensate and one fully-bleached condensate are shown both in normal mode and fire mode to enable visualization of the minimal fluorescence recovery. Mean average data are normalized to the average intensity of a particle before photobleaching and are represented as mean ± standard deviation from the recovery curves. Scale bar, 1 μm.

### The central IDR is critical for RNA-mediated phase separation

The N protein has multiple intrinsically disordered regions (IDRs) based on sequence alignments, disorder prediction, and IDR prediction algorithms including the catGRANULE server (47) (Fig. S3A): (1) the N-terminal IDR (aa 1-48), (2) the central IDR (aa 175-246) that consists of a serine/arginine rich region (aa 175-205) followed by a leucine/glutamine rich region (48) (Fig. S3C), and (3) the C-terminal IDR (aa 365-419). The folded RNA-binding N-terminal domain and the C-terminal dimerization domain are flanked by these three IDR regions (28, 49). In related betacoronavirus N proteins, each of these domains has been shown to bind RNA with different affinities (18, 43–45, 50). To determine the contribution of different domains to the protein’s ability to form phase-separated condensates with RNA, we purified a series of truncations deleting the N-terminal IDR (N^49-419^), the C-terminal IDR (N^1-364^), or both N- and C-terminal IDRs (N^49-364^). We found that while truncations removing the N- or C-terminal IDRs retained the ability to form phase-separated condensates with RNA, the resulting condensates were smaller than those formed from full-length protein, suggesting that these IDRs may play a minor role in phase separation (Fig. 3A-D). The isolated N-terminal half of the N protein (N^1-246^) underwent RNA-dependent phase sep-aration, while the isolated C-terminal half (N^247-419^) did not (Fig.3E, 3F).

**Fig. 3.**
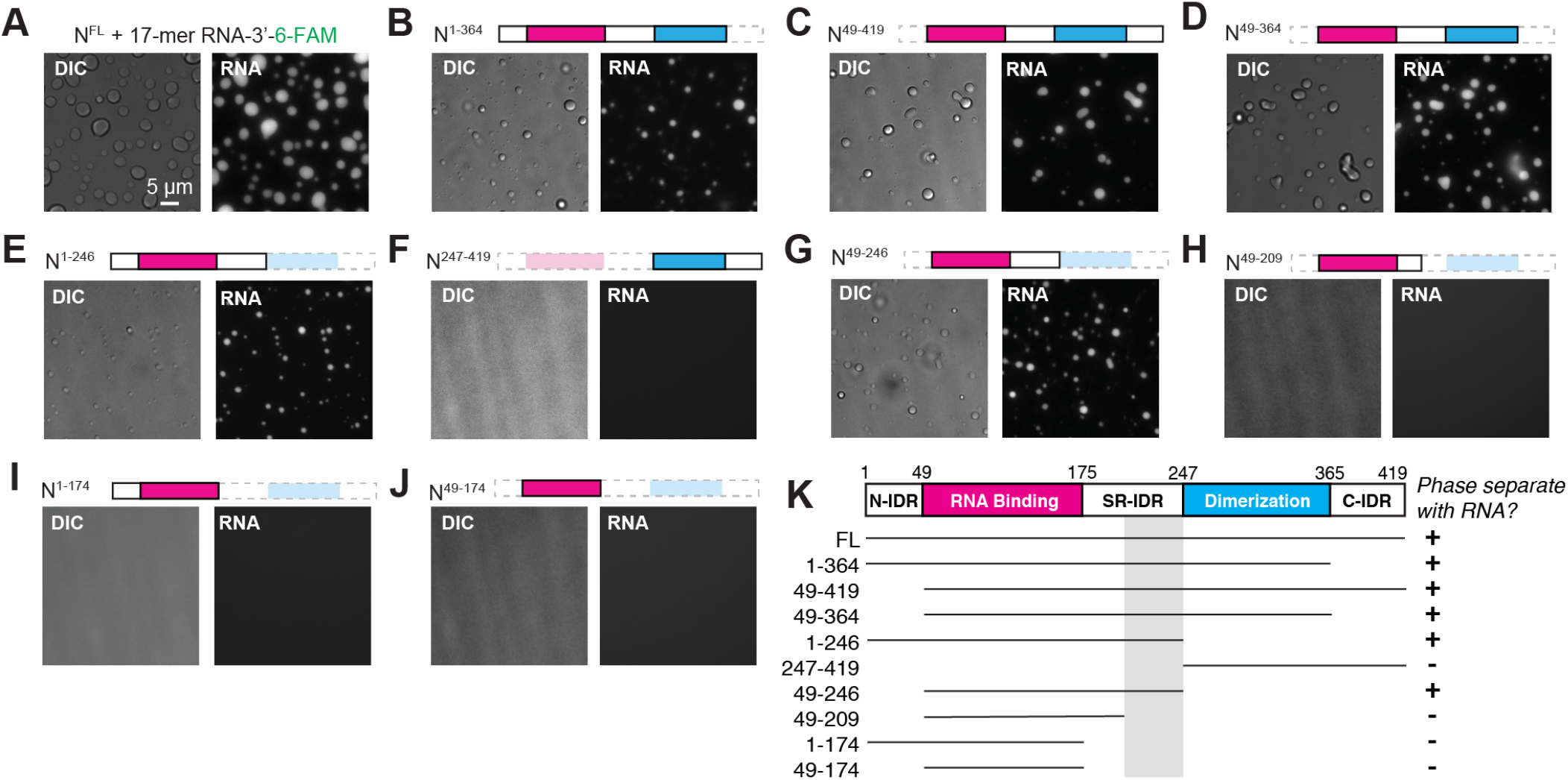
The central intrinsic disordered region is required for RNA-dependent phase separation of N protein. (A-J) DIC and fluorescence images of the mixtures of 2μM 17-mer ssRNA and 10μM N protein variants: (A) full length N^1-419^; (B) N^1-364^; (C) N^49-419^; (D) N^49-364^; (E) N^1-246^; (F) N^247-419^; (G) N^49-246^; (H) N^49-209^; (I) N^1-174^; (J) N^49-174^. See Fig. S3B for SDS-PAGE analysis of purified proteins. Images were taken within 30 min after mixing N protein variants with RNA. Scale bar, 5 μm. (K) Summary of phase separation behaviors of N protein variants shown in (A-J). Domain schematic of the SARS-CoV-2 N protein, with known domains marked. Gray shading indicates the region implicated in N+RNA phase separation.

To further delineate which domain of N^1-246^ is required for phase separation, we generated four additional truncations: N^49-246^, N^49-209^, N^1-174^, and N^49-174^. As expected, removal of the N-terminal IDR (N^49-246^) did not strongly affect RNA-mediated phase separation (Fig. 3G). As we previously found that the N^1-246^ construct is proteolytically sensitive (Fig. S3B) (21), we used mass spectrometry to map the cleavage site to residue 209 in the central IDR, just after the S/R rich subdomain. Removing residues 210-246 (N^49-209^) eliminated RNA-mediated phase separation (Fig. 3H). Constructs lacking the entire central IDR (N^1-174^ and N^49-174^) also failed to phase separate with RNA (Fig. 3I, 3J). Like the full-length protein, without RNA all these N protein truncations did not phase separate at all or at a level much lower than in the presence of RNA (Fig. S3D). Correspondingly, the truncations with an intact central IDR domain also robustly underwent phase separation with viral RNA UTR265 (Fig. S3E).

Overall, these data show that residues 210-246 in the central IDR are critical for SARS-CoV-2 N protein phase separation with RNA (Fig. 3K). While we were unable to test the effect of removing the RNA-binding NTD due to inherent instability of the isolated central IDR, we propose that RNA-induced phase separation likely requires both the RNA-binding NTD and the central IDR, with the NTD contributing most of the RNA binding capacity, and the central IDR mediating higher-order assembly and phase separation.

### The SARS-CoV-2 N protein forms liquid-like condensates in cells

In addition to its role in viral packaging, the N protein is dynamically located to the viral replicase-transcriptase complex and is required for efficient viral RNA transcription (11–15). Consistently, a recent proteomic study showed that SARS-CoV-2 N strongly interacts with cellular RNA processing machinery, including many stress granule proteins and several RNA helicases including DDX1, which has been reported to be involved in viral RNA synthesis (26, 51). These data suggest that the N protein may form phase-separated condensates in cells to promote recruitment of factors that facilitate viral RNA synthesis. To assess the ability of the N protein to phase-separate in cells, we generated a stably-transformed human osteosarcoma U2OS cell line expressing Clover-tagged SARS-CoV-2 N under the control of an inducible promoter (Fig. 4A). One day after induction of N protein expression, N protein condensates were initially formed within cells (Fig. 4A-B). Use of FRAP analysis established that the N protein was highly dynamic in these structures, with 80% fluorescence recovery within 1 min (Fig. 4B-C). This apparent liquid-like behavior strongly contrasts to N+RNA condensates that *in vitro* show a more gel-like structure, suggesting dynamic regulation of the N protein’s phase separation behavior in cells. Notably, in related coronaviruses the S/R rich subdomain in the central IDR becomes highly phosphorylated in infected cells (22, 23, 25, 26) and this subdomain’s proximity to the key region mediating phase separation suggests that condensation is likely regulated by phosphorylation. In agreement with this idea, mimicking S/R phosphorylation through serine-to-aspartic acid mutations was recently reported to increase the liquid-like properties of SARS-CoV-2 N+RNA condensates *in vitro* (52). The different behavior of phos-phorylated versus unphosphorylated forms of N may reflect adaptation to its two distinct roles, with liquid-like phosphorylated condensates promoting viral replication and gel-like unphosphorylated condensates mediate viral RNA packaging in virions (52).

**Fig. 4.**
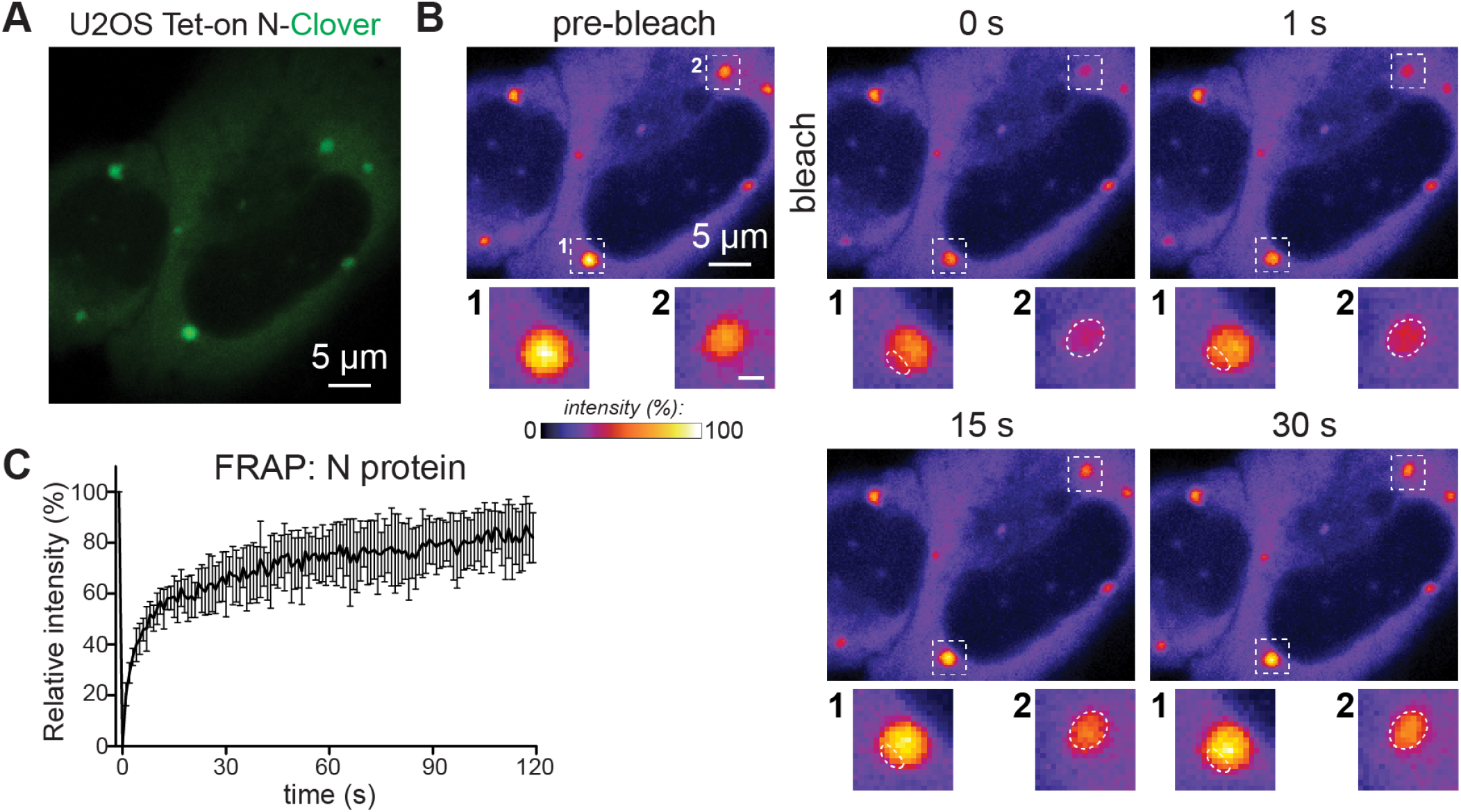
The SARS-COV-2 N protein forms a highly dynamic phase separated condensates in cells. (A) Schematic and a representative fluorescence image of N^Clover^ in U2OS cells. (B) FRAP example of the N^Clover^ condensates in the cell. Enlarged pictures are the fluorescence images of one condensate after partial photobleaching (1) and one after full photobleaching (2). Scale bar, 5 μm for original image and 1 μm for enlarged images. (C) Mean fluorescence intensity plot of N^Clover^ condensates in cell of the FRAP experiments, n =7. Mean average data are normalized to the average intensity of a particle before photobleaching and are represented as mean ± standard deviation from the recovery curves.

### The SARS-CoV-2 M protein independently promotes N protein phase separation

The abundant transmembrane M protein acts as an organizational hub for virion assembly through its binding to both the membrane-anchored spike protein and to the N protein/viral RNA complex via a soluble C-terminal domain (CTD) extending into the virion (9, 53) (Fig. 5A). In related coronaviruses, direct interactions between M and N have been reported, with the C-terminal IDR of N particularly implicated in this interaction (9, 10). To better understand how the SARS-CoV-2 M and N proteins interact to mediate virion assembly, we purified the soluble C-terminal domain of SARS-CoV-2 M (aa 104-222) fused to GFP (Fig. 5A, S4A) and mixed it with purified Cy5-labeled N protein. Purified GFP-M^104-222^ induced N protein demixing even in the absence of RNA and that the size of phase-separated condensates increased with GFP-M^104-222^ concentration (Fig. 5B, S4B). By itself, GFP-M^104-222^ did not form condensates (Fig. 5A). At higher concentration, N+M condensates formed amorphous aggregate-like structures, suggesting that M protein promotes the nucleation of N protein into dense gel-like assemblies. FRAP analysis confirmed the solid-like property of these structures (Fig. S4C).

**Fig. 5.**
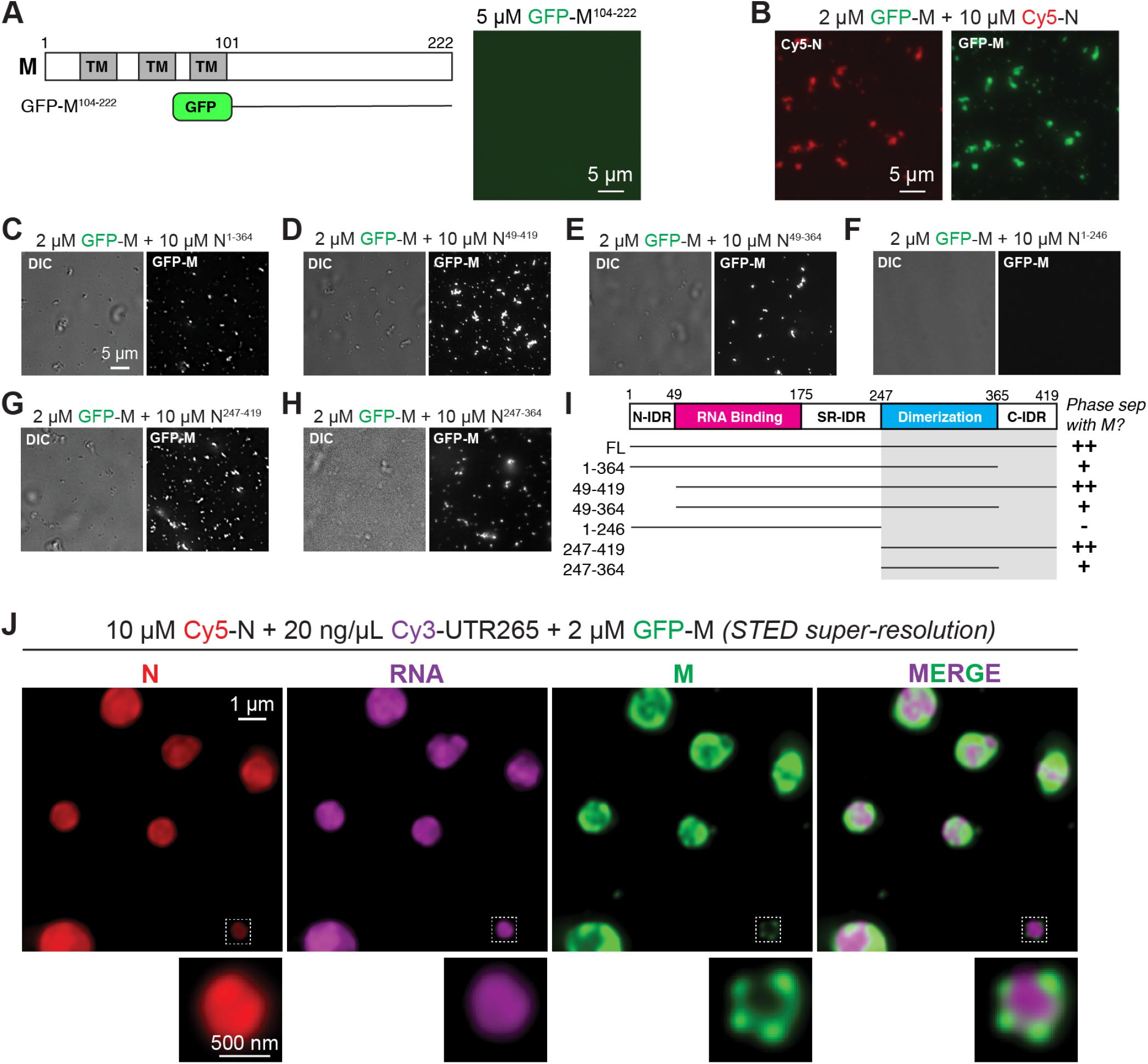
SARS-COV-2 M protein promotes N protein phase separation independent of RNA. (A) Schematic figure of the domains of M protein and the construct of GFP-M^104-222^. Fluorescence image of 5 μM GFP-M^104-222^ only. Scale bar, 5 μm. (B) Fluorescence images of phase separation of N protein when mixed with GFP-M^104-222^. N protein (10% Cy5-labeled) is used for this experiment. Scale bar, 5μm. (C-I) Representative DIC and fluorescence images of the mixtures of 2μM GFP-M^104-222^ and N protein truncation variants: (C) N^1-364^; (D) N^1-246^; (E) N^247-419^; (F) N^49-419^; (G) N^49-364^; (H) N^247-364^. Images were taken in 20 mins after mixing. Scale bar, 5μm. (I) A summary of the phase separation behavior of N protein truncation variants when mixed with 2μM GFP-M^104-222^. (J) Representative STED super-resolution images of 10μM N protein, 20 ng/μL UTR265 and 2 μM GFP-M^104-222^ were mixed subsequently within 1 min. Enlarged images represents one sub-micron condensates formed with N protein/UTR265 in the core while GFP-M^104-222^ on the outside layer. Scale bar, 1 μm for original image and 500 nm for enlarged figures.

Next, we used our truncations of the SARS-CoV-2 N protein to determine the region required for phase separation with M, focusing particularly on the previously-implicated C-terminal region (Fig. 5C-I). We found that removal of the N terminal IDR (N^49-419^) does not affect N/M de-mixing. Further removal of the C-terminal IDR (N^49-364^) decreases the level of de-mixing, suggesting that this domain plays a role in M protein binding. Removal of the C-terminal dimerization domain (N^1-246^) completely eliminated N/M phase separation (Fig. 5F), while the C-terminal part of N protein (N^247-419^), containing the C-terminal dimerization domain and C-terminal IDR, was sufficient for M protein-dependent phase separation (Fig. 5G). An N protein fragment retaining only the C-terminal dimerization domain (N^247-364^) phase separated with M, albeit to a lesser extent than N^247-419^ (Fig. 5H). RNA did not induce M protein phase separation in the absence of N (Fig. S4D). These results support the idea that the N protein C-terminal region interacts with M, and that this interaction can promote the assembly of gel-like condensates in the absence of RNA. Further, RNA-mediated and M-mediated phase separation rely on distinct domains of the N protein.

Finally, we explored the behavior of three-component systems including N, M, and RNA. Within 20 minutes of addition of GFP-M^104-222^ to pre-assembled condensates of N+RNA, GFP-M^104-222^ formed an annular shell on the condensate surface that was then stable for hours (Fig. S5A). Even more strikingly, when N, M, and RNA were all mixed simultaneously, RNA and M formed mutually exclusive condensates with N. In condensates that formed with N+M+RNA, M and RNA occupied mutually exclusive subdomains, for example, with a central core of N+RNA surrounded by a shell of N+M, or vice versa (Fig. 5J, S5B). Thus, despite the N protein forming condensates with RNA and the M protein through distinct domains, these condensates appear to be mutually exclusive.

## Discussion

A critical step in the life cycle of any virus is the packaging of its genome into new virions. This is an especially challenging problem for betacoronaviruses like SARS-CoV-2, with its large ~30 kb RNA genome. Here, we demonstrate that the SARS-COV-2 Nucleocapsid (N) protein, the structural protein largely responsible for binding, compacting, and packaging the viral genome, undergoes phase separation into gel-like condensates with the SARS-COV-2 M protein and RNA, including RNAs derived from the viral genome 5’ UTR and the region corresponding to the putative packaging signal of SARS-CoV. Our findings parallel recent reports of RNA-mediated phase separation by the nucleocapsid proteins from other viruses, including Measles virus and HIV-1 (54, 55). Our findings also parallel not-yet peer-reviewed reports of SARS-CoV-2 N+RNA phase separation (52, 56–59). These findings, plus recognition that many other viral nucleocapsid proteins possess domains with high predicted disorder (60), suggest a general role for phase separation in viral genome packaging and virion assembly. We pinpoint the region of SARS-CoV-2 N responsible for RNA-mediated phase separation as a subdomain of the protein’s central IDR, residues 210-246, which is adjacent to the phospho-regulated serine/arginine (S/R)-rich subdomain (Fig. 6A). A recent molecular dynamics analysis of this region identified a putative hydrophobic *α*-helix spanning residues ~213-225, supporting the idea that this region may be directly involved in protein-protein interactions to promote phase separation (58). We also found that a soluble fragment of the viral membrane-associated M protein interacts with N and independently induces the formation of phase-separated condensates. Three-component mixtures give rise to structures strongly reminiscent of the expected architecture of these components in virions (33, 61) with a central core of N+RNA surrounded by a shell of N+M.

**Fig. 6.**
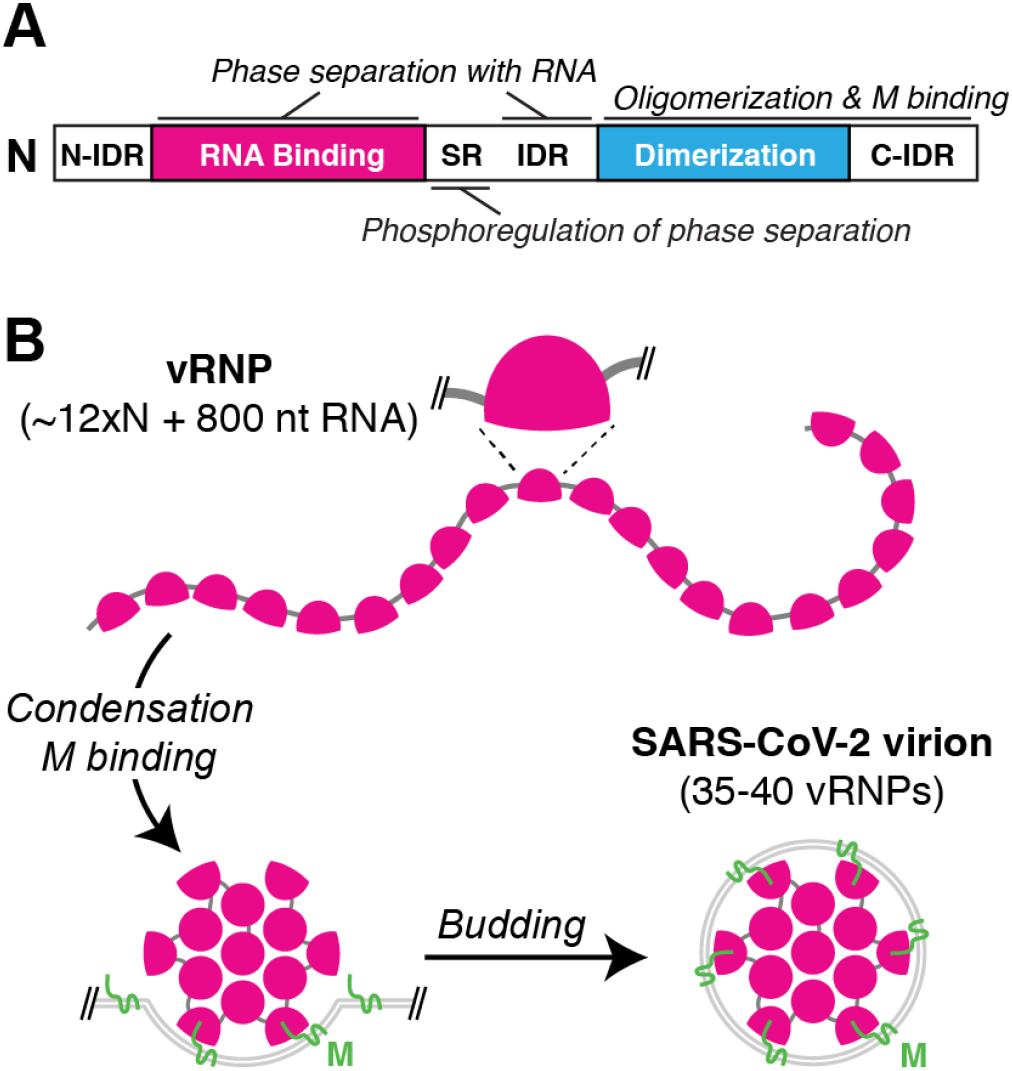
Model for SARS-CoV-2 virion assembly. (A) Domain organization of SARS-CoV-2 N protein, with domains responsible for phase separation with RNA (RNA-binding domain and central IDR) and M protein (C-terminal domains) marked. Related work suggests a role for phosphorylation of the SR-rich region in regulating the gel vs. liquid behavior of N+RNA condensates (52). (B) Proposed assembly mechanism of SARS-CoV-2 virions. Viral RNPs with ~12 copies of N and 800 nt RNA33 coat the genomic RNA, and RNA-driven condensation plus interactions with M drive RNA packaging into virions. Within the virion, individual vRNPs show characteristic orientation on the viral envelope (33) supporting a role for specific N+M interactions in packaging.

Our data on N-RNA and N-M interactions have important implications for understanding how the SARS-CoV-2 genome is packaged into developing virions. Recent high-resolution cryo-electron tomography of intact SARS-CoV-2 virions has revealed that individual viral RNA-protein complexes (vRNPs) adopt a characteristic shell-like architecture ~15 nm in diameter, comprising ~12 N proteins and --’-800 nt of RNA (4, 33). Negative-stain electron microscopy of soluble N+RNA complexes has revealed similar particles (52), suggesting that viral RNA packaging is mediated first by assembly of individual vRNPs along the genomic RNA, followed by condensation of these RNPs and recruitment to developing virions through interactions with the M protein at the cytoplasmic side of the ER-Golgi intermediate compartment (Fig. 6B) (4, 61). Our finding that the M protein can independently mediate phase separation of N suggests that a network of membrane-associated M proteins could mediate recruitment of N+RNA condensates into the developing virion (61).

Given the variety of RNAs in infected cells, including both host mRNAs and viral subgenomic RNAs, how is the full-length viral genome specifically recruited and packaged into virions? Current models posit that the N protein likely binds specifically to a viral RNA sequence that adopts a characteristic 3D structure. A recent computational analysis offers a compelling model for how phase separation might contribute to packaging specificity. Specifically, high-affinity binding of N to a particular site in the 30 kb genome could seed the assembly of vRNPs along the RNA through lower-affinity interactions, and drive condensation of stable “single-genome condensates” that can then be packaged into a virion (58). This model is indirectly supported by the recent observation of dense virus-like particles ~50-80 nm in diameter in infected cells, which may represent N+RNA condensates that have not yet been enveloped (33). The specific packaging signal is not identified for SARS-CoV-2 but may be located in the 5’ UTR or within the ORF1ab gene (43, 44, 56). We find that N forms condensates with diverse RNAs, but that the morphology of condensates formed with the sequence equivalent to the putative SARS-CoV packaging signal (PS576) is distinct from those formed with other RNAs. While this observation suggests that specific RNAs could alter the phase separation behavior of N, parallel studies have shown that altering the RNA-protein ratio in condensates can give rise to similar morphological changes (52). Further study will be required to both identify the SARS-CoV-2 packaging signal and determine how it modulates N+RNA phase separation to promote packaging of single genomes into developing virions.

In addition to their central role in viral RNA packaging, betacoronavirus N proteins participate in viral RNA transcription, especially the discontinuous transcription of subgenomic mRNAs (62). In cells, the protein is dynamically recruited to replicase-transcriptase complexes through an interaction with viral replicase subunit NSP3 (12, 63). This activity is promoted by hyperphosphorylation of the N protein in the S/R-rich region within the central IDR, and phosphorylated N protein can recruit host factors like RNA helicases to promote viral RNA template switching and subgenomic mRNA transcription (26). In contrast, N protein incorporated into virions is hypophosphorylated (25, 26). We and others have shown that N protein condensates in cells show a more liquid-like behavior than those assembled *in vitro*, and recent reports directly implicate phosphorylation of the S/R-rich region in this behavior difference (52, 56). Given the proximity of the S/R-rich region (aa 175-205) to the region we implicate in RNA-mediated phase separation (aa 210-246), it is not surprising that phosphorylation in this region could strongly affect N-protein self-interactions and alter the properties of the resulting condensates. In this manner, the physical properties of N+RNA condensates could be tuned depending on whether it is promoting viral transcription (hyperphosphorylated, liquid-like) or RNA packaging (hypophosphorylated, gel-like).

The ongoing COVID-19 pandemic demands a concerted effort to develop new therapeutic strategies to prevent new infections and lessen the severity of the infection in patients. The concurrent finding by a number of groups (52, 56–59) that the SARS-CoV-2 N protein undergoes phase separation with RNA, and that this behavior is likely critically involved in viral RNA packaging during virion development, offers a new step in the viral life cycle that could be targeted with therapeutics. Our finding that phase separation also drives interactions with the viral M protein highlights another interaction that could be targeted to disrupt the viral life cycle. Our observations that purified N protein readily phase-separates with RNA or M protein *in vitro*, and that N forms condensates when expressed in mammalian cells, suggest several straightforward experimental strategies to test whether FDA-approved drugs or novel therapeutics may disrupt these assemblies. Such a strategy could provide an important complement to current therapeutic efforts targeting other steps in the viral life cycle.

## Materials and Methods

### Cloning and Protein Purification

SARS-CoV-2 N protein constructs (N^FL^, N^1-364^, N^49-419^, N^49-364^, N^1-264^, N^247-419^, N^49-246^, N^49-209^, N^1-174^, N^49-174^) were amplified by PCR from the IDT 2019-nCoV N positive control plasmid (IDT cat. # 10006625; NCBI RefSeq YP_009724397) and inserted by ligation-independent cloning into UC Berkeley Macrolab vector 2B-T (AmpR, N-terminal His_6_-fusion; Addgene #29666) for expression in *E. coli.* The gene expressing M protein residues 104-222 was synthesized by IDT, and inserted into UC Berkeley Macrolab vector 1GFP (KanR, N-terminal His_6_-GFP fusion; Addgene #29663). Plasmids were transformed into *E. coli* strain Rosetta 2(DE3) pLysS (Novagen), and grown in the presence of ampicillin or kanamycin and chloramphenical to an OD_6_00 of 0.8 at 37°C, induced with 0.25 mM IPTG, then grown for a further 16 hours at 18°C prior to harvesting by centrifugation. Harvested cells were resuspended in buffer A (25 mM Tris-HCl pH 7.5, 5 mM MgCl_2_, 10% glycerol, 5 mM *β*-mercaptoethanol, 1 mM NaN3) plus 1 M NaCl (N^FL^, N^1-364^, N^49-419^, N^49-364^) or 500 mM NaCl (all other constructs) and 5 mM imidazole pH 8.0.

For purification, cells were lysed by sonication, then clarified lysates were loaded onto a Ni^2+^ affinity column (Ni-NTA Superflow; Qiagen), washed in buffer A plus 300 mM NaCl and 20 mM imidazole pH 8.0, and eluted in buffer A plus 300 mM NaCl and 400 mM imidazole. For cleavage of His_6_-tags, proteins were buffer exchanged in centrifugal concentrators (Amicon Ultra, EMD Millipore) to buffer A plus 300 mM NaCl and 20 mM imidazole, then incubated 16 hours at 4°C with TEV protease. Cleavage reactions were passed through a Ni^2+^ affinity column again to remove uncleaved protein, cleaved His_6_-tags, and His_6_-tagged TEV protease. Proteins were concentrated in centrifugal concentrators and purified by size-exclusion chromatography (Superpose 6 Increase 10/300 GL or Superdex 200 Increase 10/300 GL; GE Healthcare) in gel filtration buffer (25 mM Tris-HCl pH 7.5, 300 mM NaCl, 5 mM MgCl_2_, 10% glycerol, 1 mM DTT). Purified proteins were concentrated and stored at 4°C for analysis.

For fluorescent labeling, we inserted a cysteine residue between the N-terminal His_6_-tag and the N-terminal residue of the full-length N protein, and purified the protein as above. We labeled with either Alexa Fluor 488 (Thermo Fisher) or sulfo-Cyanine 5 (Lumiprobe) according to Lumiprobe’s protocol “Maleimide labeling of proteins and other thiolated biomolecules”. Briefly, an excess of TCEP (tris-carboxyethylphosphine, up to 100x molar) was added to protein between 1-10 mg/mL, and kept the mixture for 20 minutes at room temperature, then added 1/20 x fold dye solution, mixed well and left overnight at 4°C. Next day, proteins were separated from unreacted fluorophores by passing over a Superpose 6 Increase 10/300 GL column (GE Healthcare).

### *In vitro* transcription

DNA templates for viral RNA fragments were ordered as gBlocks from IDT and amplified by PCR through primers with T7 promoter sequence (TAAT-ACGACTCACTATAGGGAG) in the 5’ end. Genomic RNA fragments (Table S1) were synthesized following the protocol of HiScribe T7 high yield RNA synthesis kit (NEB, E2040S) with 1 μg purified PCR products. To label RNA, 0.1 μL of 10 mM Cy3-UTP (Enzolifesciences, 42506) was added to the transcription system. After DNase I treatment, RNAs were purified using Trizol/Chloroform isolation method and solubilized with 5 mM HEPES, pH 7.5 RNase-free buffer. The size and purity of RNAs were verified by denaturing Urea/PAGE, and their concentrations measured by Nanodrop.

### In *vitro* phase separation assays

*In vitro* phase separation assays are performed at 20°C unless otherwise indicated. N protein mixed with Cy5-labeled or Alexa-488 labeled protein at 1:100 or 1:10 ratio was used for phase separation assay. Phase separation of N protein (in 5 mM HEPES, pH 7.5, 80 mM KCl) was induced by either adding of RNA (in 5 mM HEPES, pH 7.5) or M protein (in 5 mM HEPES, pH 7.5, 80 mM KCl). Samples were mixed in protein LoBind tubes (Eppendorf, 022431064) and then immediately transferred onto 18-well glass bottom chamber slide (iBidi, 81817). Condensates were imaged within 10-20 mins or otherwise as indicated in the experiment.

### Turbidity assay

Samples were mixed in protein LoBind tubes as in the previous section. After 30 mins, the turbidity of each sample was measured by nanodrop at 350 nm absorbance.

### Stable human cell line construction

For expression in U2OS cells, the SARS-CoV-2 N protein was cloned into a third-generation Tet-on system on a lentivirus vector with a Clover tag at the C-terminal of N protein. Lentivirus is produced in HEK293t cells by transfection of lentiviral plasmid and packaging plasmids pMD2.G and psPAX2. After two days of transfection, the culture medium containing the lentivirus was passed through a 0.45 μm filter and was used to infect U2OS cell line. After two days of infection, the medium is exchanged to medium containing 20 μg/μL blasticidin for selection.

### Microscopy of *in vitro* phase separated condensates

DIC and fluorescence images of *in vitro* condensates were taken on a DeltaVision Elite (GE Healthcare) microscope with a 60x oil N.A. 1.42 objective. The laser power and exposure time of each channel are optimized to avoid saturation. Alternatively, condensates were imaged using Zeiss LSM880 or Leica SP8 STED superresolution confocal microscopes. Laser power for imaging, digital offset and gain values are optimized for each channel to ensure the intensity lies in the linear range. For STED superresolution imaging, image resolution settings are optimized to reach ~20 nm/pixels; average of the intensity from 16 images to increase the signal to noise ratio; 50% of 775 nm STED depletion laser is used for STED imaging. Time lapse imaging was used to capture fusion of RNA/N protein condensates on a DeltaVision microscope at 1 min intervals.

### Fluorescence recovery after photobleaching (FRAP)

FRAP analysis of condensates *in vitro* was performed on a Zeiss LSM880 Aryscan microscope with 63x/1.42 oil objective or 40x/1.2 W objective or on Nikon Eclipse Ti2 A1 confocal microscope as indicated. The intensity of the fluorescent signal is controlled in the detection range through changing the laser power, digital gain and off-set. For green, red and far-red fluorescent channels, bleaching was conducted by 488-nm, 561-nm or 633-nm line correspondingly and the laser power and iteration of bleaching are optimized to get an efficient bleaching effect. Fluorescence recovery was monitored at 2 s or 4 s intervals for 4 min. In the focal-bleach experiment, roughly half (partial bleach) or all (full bleach) of a condensate was photobleached to determine the molecular mobility with diffuse pool or inside a condensate. U2OS cells for FRAP experiments were cultured on a 8-well chamber slide (iBidi, 80827) in DMEM supplemented with 10% fetal bovine serum (FBS) and Antibiotic-Antimycotic (Ther-mofisher, 15240062). N^Clover^ expression was induced for 24 hours by adding 1 μg/mL doxycycline to the culture medium. FRAP experiment was conducted as for *in vitro* condensate analysis, using a Zeiss LSM88microscope. The FRAP data were quantified using ImageJ or Zeiss Zen built-in profile model. The time series of the fluorescence intensity of condensates were calculated. The intensity of the condensate during the whole experiment was normalized to the one before bleaching and the intensity of the granule just after bleaching was normalized to zero. At least 2-10 condensates per condition were analyzed to calculate the mean and standard deviation. The averaged relative intensity and standard error were plotted to calculate dynamics.

### Cryo-electron tomography

For cryo-electron tomography, phase separation was induced by mixing 10 μM N protein (in 5 mM HEPES, pH 7.5, 80 mM KCl) with 20 ng/μL of the indicated RNA (in 5 mM HEPES, pH 7.5) for 0.5-1 minute at 20°C. 4μL of the resulting solution was deposited on glow-discharged Quantifoil grids (R2/1, Cu 200-mesh grid, Electron Microscopy Sciences). The solution on the grid was then blotted using a Vitrobot (Thermo Fisher Scientific) with conditions set to blot force −10, blot time 2.5-3.5 seconds, and drain time 2 seconds. After blotting, the grid was immediately plunge-frozen into liquid ethane/propane mixture (Airgas) cooled to close to liquid nitrogen temperature. Cryo-electron tomography (cryo-ET) imaging was performed on a Titan Krios operated at 300 KeV (Thermo Fisher Scientific) equipped with a post-column Quantum energy filter (Gatan). The images were recorded on a K2 Summit direct detector (Gatan) in counting mode using SerialEM (64). The tilt-series were acquired using a dose-symmetric scheme (65) with a tilt range of ± 60° in EFTEM mode with a nominal magnification of 42,000x-64,000x (pixel sizes at the camera 0.34-0.22 nm), tilt increment: 2-3°, and a target defocus of −3-5 μm. Motion-correction and dose-weighting, using MotionCor2 (66), was applied to the individual frames of the tilt-series images. The motion-corrected and dose-weighted tilt-series images were then aligned in IMOD (67) using patch-tracking. 3D CTF correction was performed using novaCTF (68). The weighted back-projection method was used for the final tomographic reconstruction.

## AUTHOR CONTRIBUTIONS

SL conceived of the project and planned the experiments with advice from KDC and DWC. SL performed all microscopy experiments, QY purified all proteins, and DS performed cryo-electron tomography experiments. All authors interpreted data. SL and KDC prepared figures and wrote the manuscript with input from QY, DS, EV, and DWC. EV, KDC, and DWC obtained funding.

## ACKNOWLEDGEMENTS

We thank Jennifer Santini at the UCSD Microscopy Core and Eric Griffis at the UC San Diego Nikon Imaging Center for assistance with imaging, and Huilin Zhou for assistance with mass spectrometry. We are grateful for helpful discussions and suggestions from Michael Baughn, Prasad Trivedi, Haiyang Yu, Sonia Vazquez-Sanchez and Pablo Lara Gonzalez. DS is supported by the Damon Runyon Cancer Research Foundation (DRG-2364-19). EV acknowledges funding from the NIH (DP2 GM123494), KDC acknowledges institutional funding from UC San Diego, and DWC acknowledges support from the NIH (R01 NS27036) and the Nomis Foundation. We acknowledge instrumentation support for mass spectrometry from NIH S10 OD023498. We acknowledge the use of the UC San Diego cryo-EM facility, which was built and equipped with funds from UC San Diego and an initial gift from Agouron Institute. We thank the Henriques Lab for the bioRxiv LATEX template.

