## Supplemental Material for "The SARS-CoV-2 Nucleocapsid phosphoprotein forms mutually exclusive condensates with RNA and the membrane-associated M protein"

**Figures S1-S5**
**Table S1**

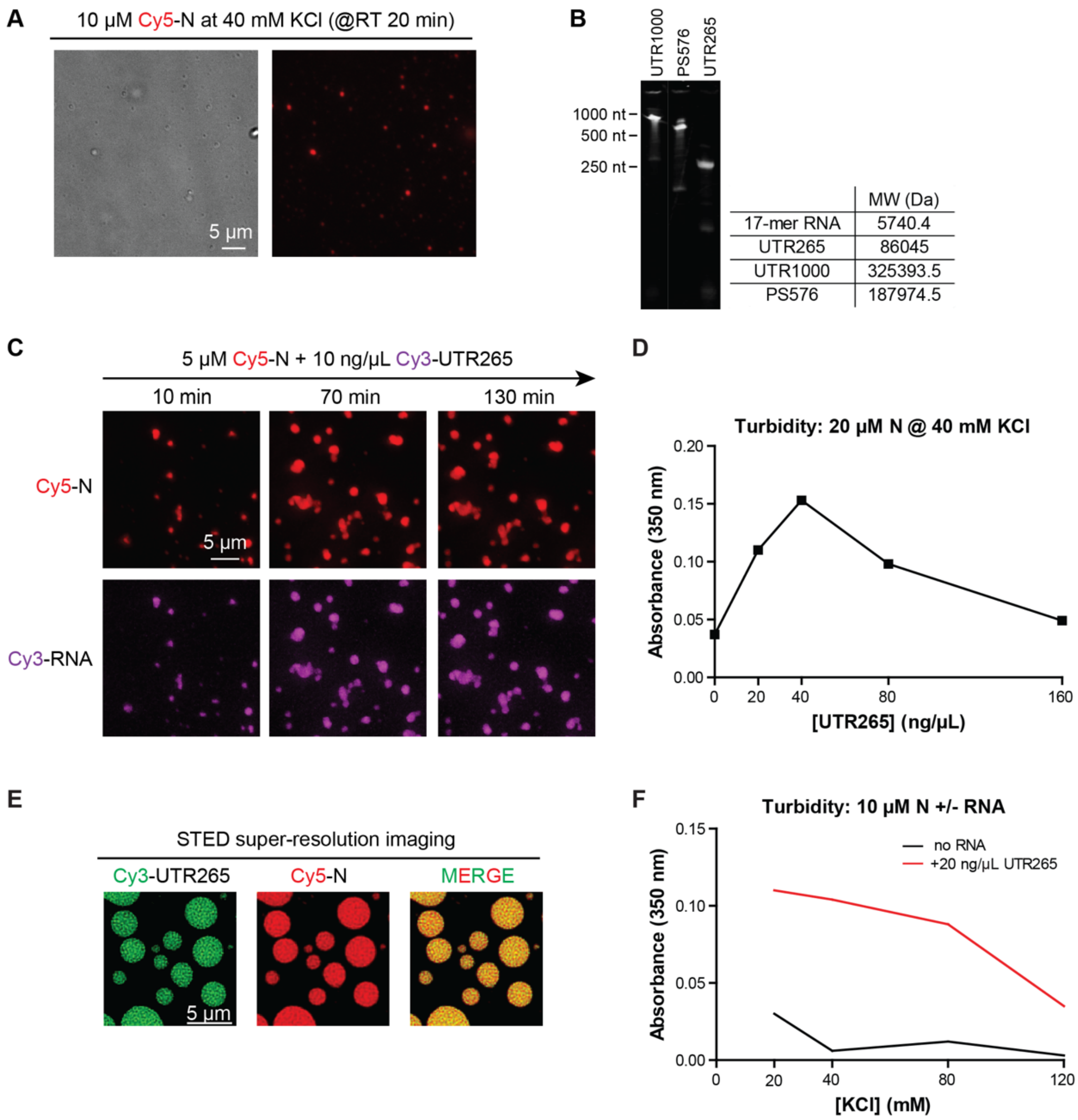


| | MW (Da) |
|---|---|
| 17-mer RNA | 5740.4 |
| UTR265 | 86045 |
| UTR1000 | 325393.5 |
| PS576 | 187974.5 |

**Fig. S1.** Characterization of N protein+RNA condensates. (A) Example of condensate formation with isolated N protein. While the protein was isolated in high-salt buffers (1M NaCl) and showed the expected $A_{260}/A_{280}$ ratio for pure protein, this phase separation behavior could be attributable to residual contaminating bacterial nucleic acids. Scale bar, 1 $\mu$m. (B) Analysis of in vitro transcribed viral RNA fragments by denaturing Urea-PAGE. See **Table S1** for sequences. (C) Time lapse imaging of N protein + UTR265 condensates. Scale bar, 5 $\mu$m. (D) Turbidity analysis of N+RNA (UTR265) mixtures at 40 mM KCl and different RNA concentrations. (E) STED super-resolution image of N+UTR265 condensates. Scale bar, 5 $\mu$m. (F) Turbidity analysis of N+RNA (UTR265) mixtures at different salt concentrations.

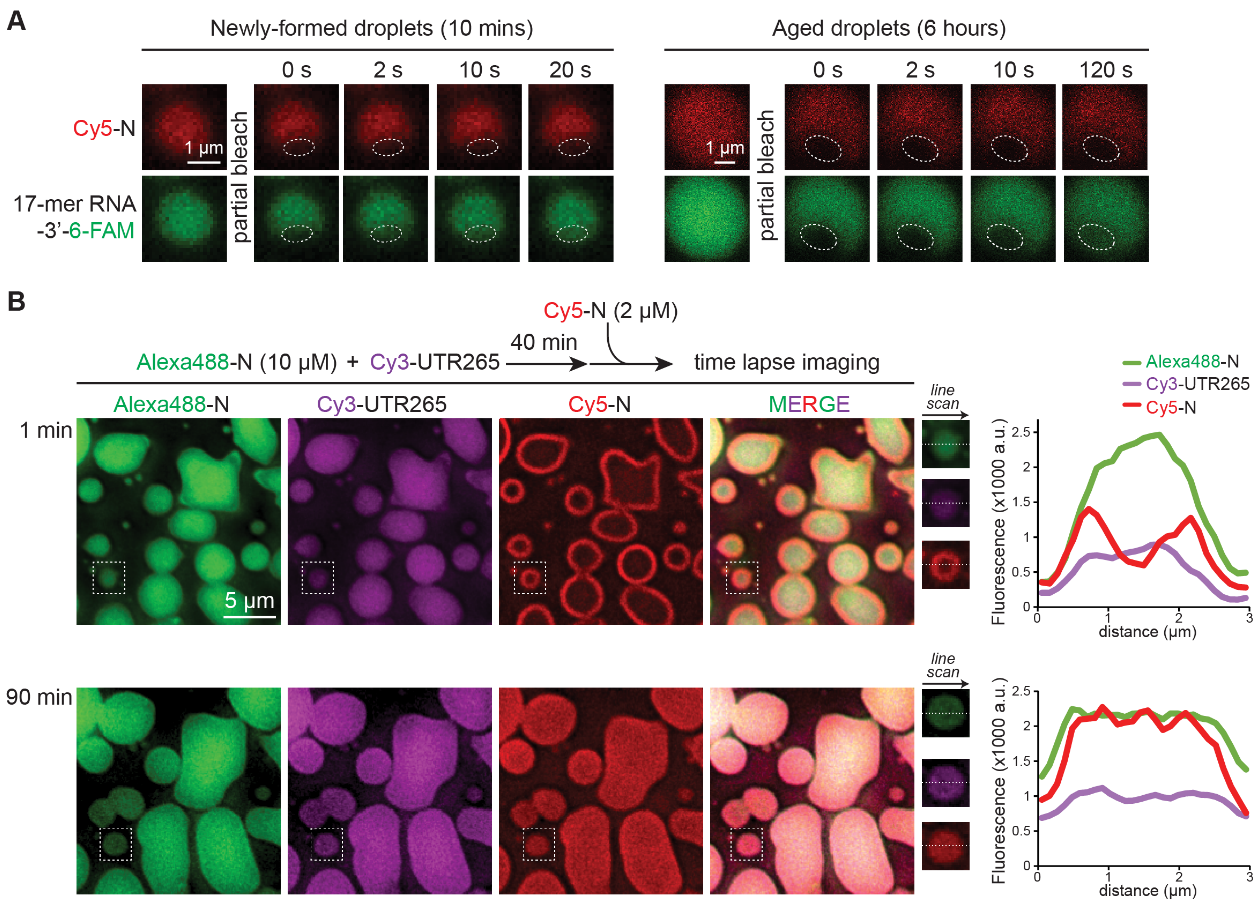


**Fig. S2.** Characterization of N+RNA condensates. (A) Fluorescence images of partial FRAP assay of newly formed and aged 17-mer N+RNA condensates. Scale bar, 1 $\mu$m. (B) Fluorescence images of the process of newly added N proteins penetration into N+UTR265 protein condensates. N+UTR265 protein (10% Alexa488-labeled) condensates were preassembled for 40 min before adding 2 $\mu$M additions N protein (10% Cy5-labeled). The enlarged picture represents the localization of Alex488-N, UTR265, and Cy5-N in one droplet, 1 min or 90 mins after adding Cy5-N protein. *Right:* Fluorescence intensity plot Alexa488, Cy3 and Cy5 channels from line scans across the indicated droplet at 1 min and 90 mins.

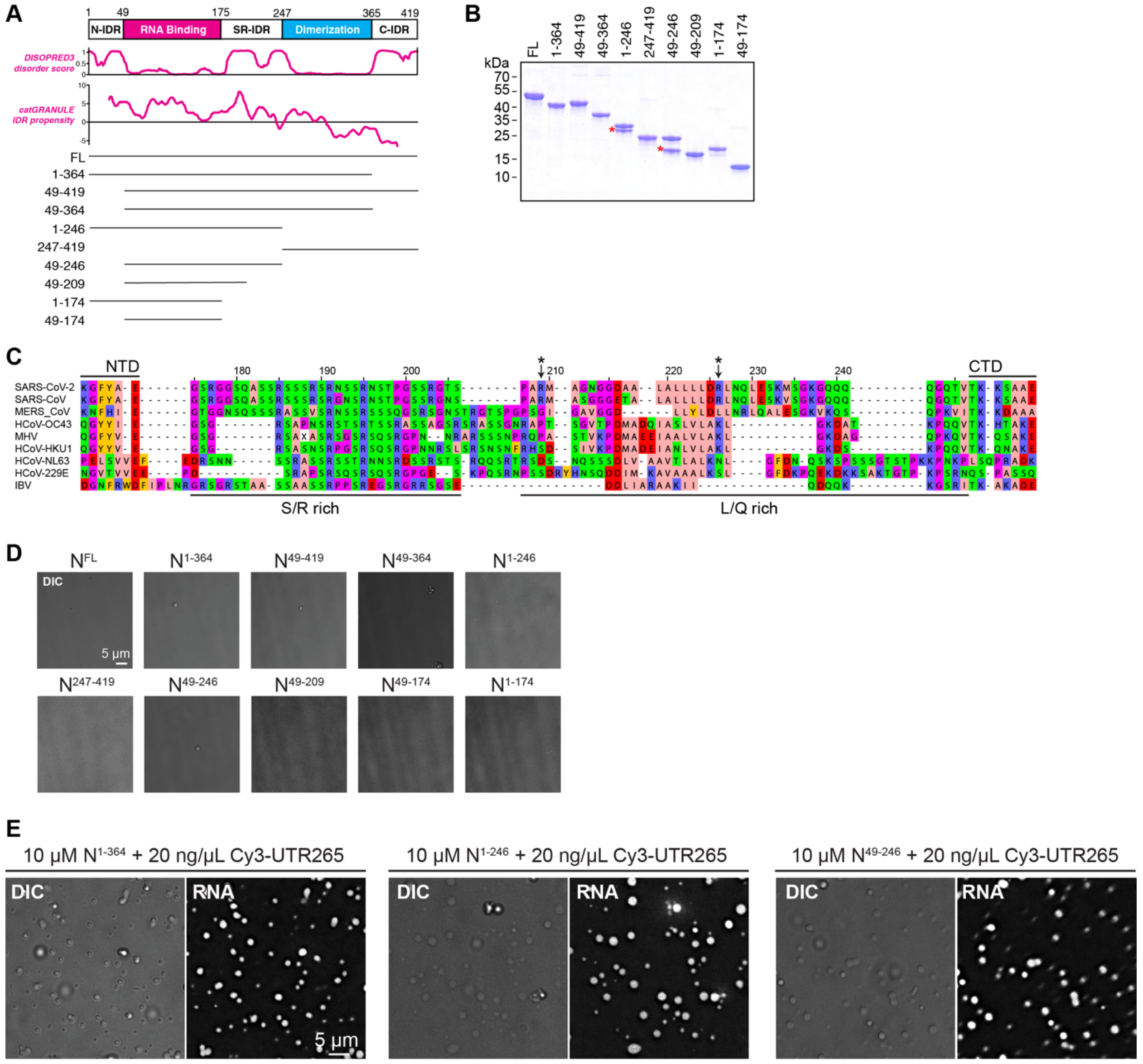


**Fig. S3.** Determination of domains involved in N+RNA phase separation. (A) N protein domains aligned with disorder propensity calculated by the DISOPRED3[48] server and catgranule[47] IDR propensity score analysis. (B) SDS-PAGE analysis of all the N protein variants that are used for in vitro phase separation assay in **Fig. 3**. (C) Sequence alignment of nine related coronavirus N proteins (SARS-CoV-2 NCBI Refseq ID QJA17760; SARS-CoV AYV99827; MERS-CoV QBM11755; HCoV-OC43 QBP84763; MHV AWB14620; HCoV-HKU1 ABG77571; HCoV-NL63 ABI20791; HCoV-229E AAA45463; IBV AAB24054), showing the central IDR region. Asterisks indicate proteolytically sensitive sites identified by mass spectrometry. (D) DIC images of isolated N protein constructs. (E) DIC and fluorescence images of phase separation behavior of N protein truncations $N^{1\text{-}364}$, $N^{1\text{-}246}$, $N^{49\text{-}246}$ when mixed with UTR265. Scale bar, 5 $\mu$m.

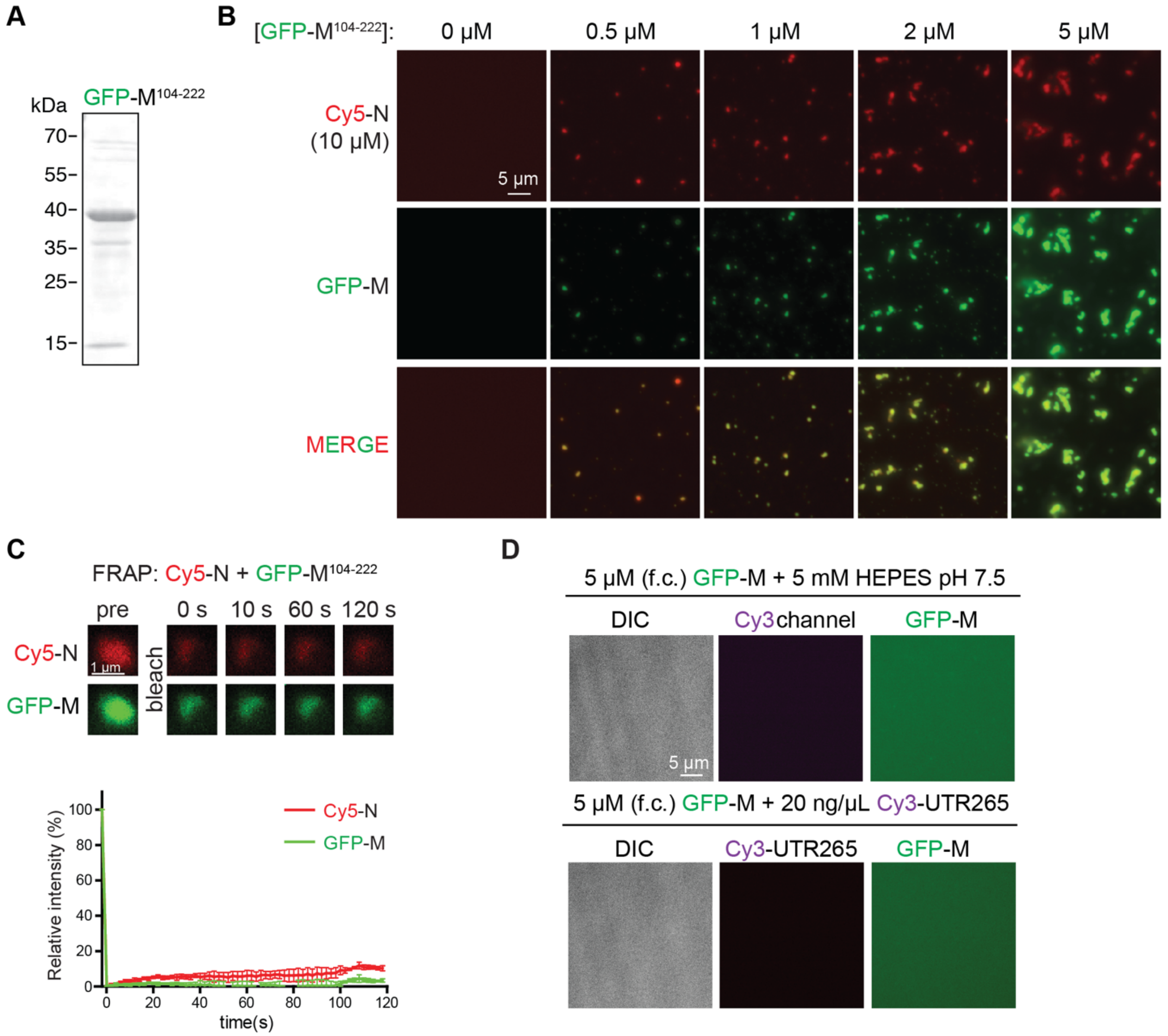


**Fig. S4.** Characterization of N+M condensates. (A) SDS-PAGE analysis of GFP-M$^{104\text{-}222}$. (B) Fluorescence images of phase separation of N protein when mixed with different concentration of GFP-M$^{104\text{-}222}$. N protein (10% Cy5-labeled) is used for this experiment. Scale bar, 5 $\mu$m. (C) *Top:* Representative images of a partial FRAP of N/M protein condensates. *Bottom:* Mean fluorescence intensity plot of N/M condensates in the FRAP experiment, n = 8. Mean average data are normalized to the average intensity of a particle before photobleaching and are represented as mean ± standard deviation from the recovery curves. (D) Representative DIC and fluorescence images of GFP-M$^{104\text{-}222}$ when mixed with 20 ng/$\mu$L Cy3-UTR265 (top) or buffer (bottom). Images were taken 20 minutes after mixing. Scale bar, 5 $\mu$m.

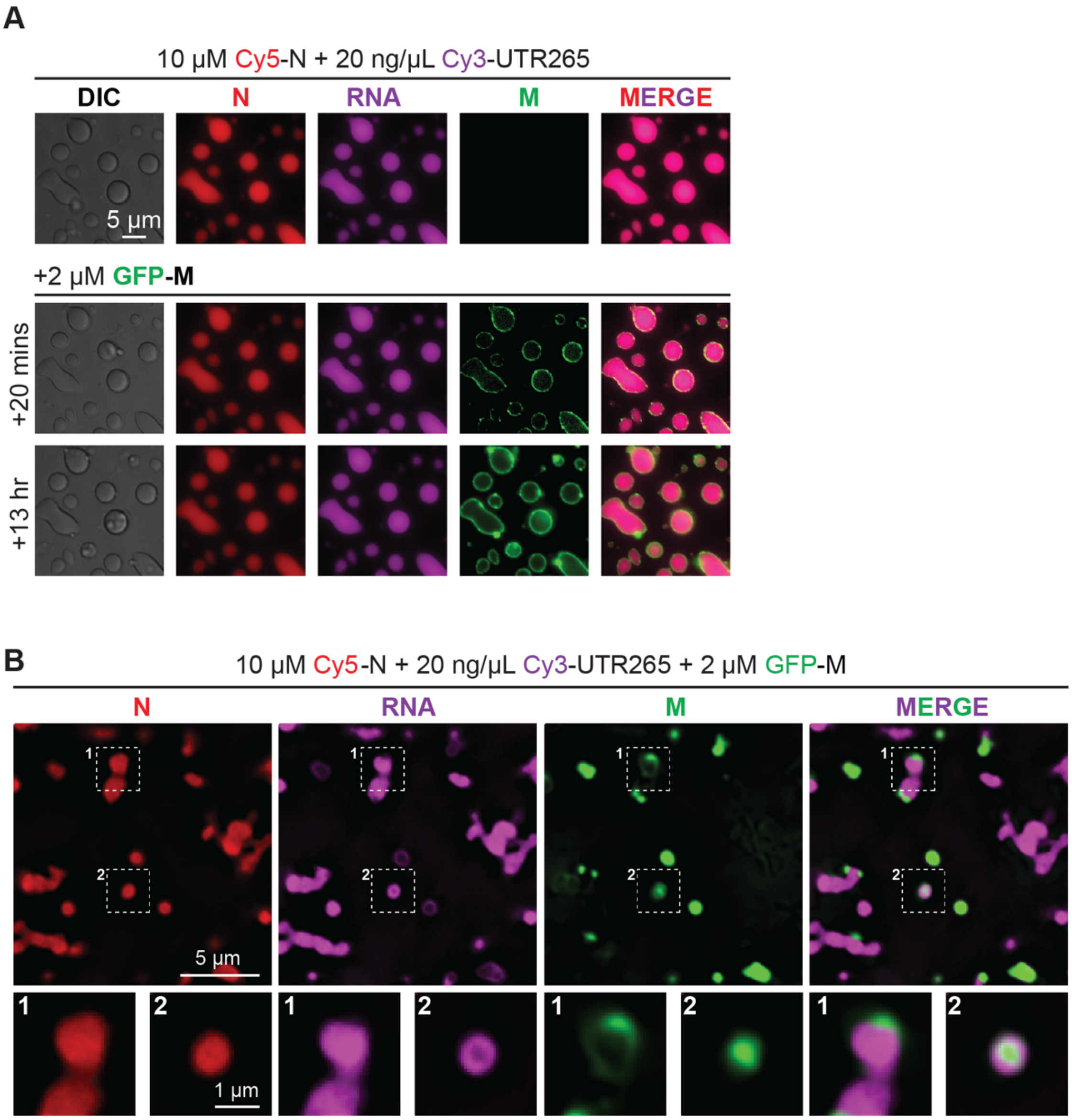


**Fig. S5.** M protein and RNA form mutually exclusive condensates with N protein. (A) Representative DIC and fluorescence images of GFP-$M^{104\text{-}222}$ when mixed with 20 ng/$\mu$L Cy3-UTR265 (top) or buffer (bottom). Images were taken in 20 minutes after mixing. Scale bar, 5 $\mu$m. M protein and RNA form mutually exclusive condensates with N protein. (A) Representative images of N+UTR265 condensates when mixed with GFP-$M^{104\text{-}222}$. Pre-assembled N+RNA condensates were formed for four hours and then 2 $\mu$M GFP-$M^{104\text{-}222}$ was added to the samples. Images were taken before and 20 min or 13 hours after adding M protein. Scale bar, 5 $\mu$m. (B) Representative images of 10 $\mu$M N protein, 20 ng/$\mu$L UTR265 and 2 $\mu$M GFP-$M^{104\text{-}222}$ were mixed subsequently within 10 min. Enlarged images are example of two circumstances: 1. M protein forming a layer on the surface of N+UTR265 condensates; 2. RNA forming a layer on the surface of N+M condensates. Scale bar, 5 $\mu$m for original images and 1 $\mu$m for enlarged images.

**Table S1**. Sequences of RNAs used in this study.

All sequences derived from NCBI RefSeq NC_045512)

```
>17-mer RNA
AAGCAGCUAAGAGCGAA

>UTR265 (nt 1-265)
AUUAAAGGUUUAUACCUUCCCAGGUAACAAACCAACCAACUUUCGAUCUCUUGUAGAUCUGUUCUCUAAACGAACUUUAAAAU
CUGUGUGGCUGUCACUCGGCUGCAUGCUUAGUGCACUCACGCAGUAUAAUUAAUAACUAAUUACUGUCGUUGACAGGACACGA
GUAACUCGUCUAUCUUCUGCAGGCUGCUUACGGUUUCGUCCGUGUUGCAGCCGAUCAUCAGCACAUCUAGGUUUCGUCCGGGU
GUGACCGAAAGGUAAG

>UTR1000 (nt 1-1000)
AUUAAAGGUUUAUACCUUCCCAGGUAACAAACCAACCAACUUUCGAUCUCUUGUAGAUCUGUUCUCUAAACGAACUUUAAAAU
CUGUGUGGCUGUCACUCGGCUGCAUGCUUAGUGCACUCACGCAGUAUAAUUAAUAACUAAUUACUGUCGUUGACAGGACACGA
GUAACUCGUCUAUCUUCUGCAGGCUGCUUACGGUUUCGUCCGUGUUGCAGCCGAUCAUCAGCACAUCUAGGUUUCGUCCGGGU
GUGACCGAAAGGUAAGAUGGAGAGCCUUGUCCCUGGUUUCAACGAGAAAACACACGUCCAACUCAGUUUGCCUGUUUUACAGG
UUCGCGACGUGCUCGUACGUGGCUUUGGAGACUCCGUGGAGGAGGUCUUAUCAGAGGCACGUCAACAUCUUAAAGAUGGCACU
UGUGGCUUAGUAGAAGUUGAAAAAGGCGUUUUGCCUCAACUUGAACAGCCCUAUGUGUUCAUCAAACGUUCGGAUGCUCGAAC
UGCACCUCAUGGUCAUGUUAUGGUUGAGCUGGUAGCAGAACUCGAAGGCAUUCAGUACGGUCGUAGUGGUGAGACACUUGGUG
UCCUUGUCCCUCAUGUGGGCGAAAUACCAGUGGCUUACCGCAAGGUUCUUCUUCGUAAGAACGGUAAUAAAGGAGCUGGUGGC
CAUAGUUACGGCGCCGAUCUAAAGUCAUUUGACUUAGGCGACGAGCUUGGCACUGAUCCUUAUGAAGAUUUUCAAGAAAACUG
GAACACUAAACAUAGCAGUGGUGUUACCCGUGAACUCAUGCGUGAGCUUAACGGAGGGGCAUACACUCGCUAUGUCGAUAACA
ACUUCUGUGGCCCUGAUGGCUACCCUCUUGAGUGCAUUAAAGACCUUCUAGCACGUGCUGGUAAAGCUUCAUGCACUUUGUCC
GAACAACUGGACUUUAUUGACACUAAGAGGGGUGUAUACUGCUGCCGUGAACAUGAGCAUGAAAUUGCUUGGUACACGGAACG
UUCU

>PS576 (nt 19786-20361)
GAGCUUUGGGCUAAGCGCAACAUUAAACCAGUACCAGAGGUGAAAAUACUCAAUAAUUUGGGUGUGGACAUUGCUGCUAAUAC
UGUGAUCUGGGACUACAAAAGAGAUGCUCCAGCACAUAUAUCUACUAUUGGUGUUUGUUCUAUGACUGACAUAGCCAAGAAAC
CAACUGAAACGAUUUGUGCACCACUCACUGUCUUUUUUGAUGGUAGAGUUGAUGGUCAAGUAGACUUAUUUAGAAAUGCCCGU
AAUGGUGUUCUUAUUACAGAAGGUAGUGUUAAAGGUUUACAACCAUCUGUAGGUCCCAAACAAGCUAGUCUUAAUGGAGUCAC
AUUAAUUGGAGAAGCCGUAAAAACACAGUUCAAUUAUUAUAAGAAAGUUGAUGGUGUUGUCCAACAAUUACCUGAAACUUACU
UUACUCAGAGUAGAAAUUUACAAGAAUUUAAACCCAGGAGUCAAAUGGAAAUUGAUUUCUUAGAAUUAGCUAUGGAUGAAUUC
AUUGAACGGUAUAAAUUAGAAGGCUAUGCCUUCGAACAUAUCGUUUAUGGAGAUUUUAGUCAUAGUCAGUUAGGUGGU
```
